# Large-scale functional integration, rather than functional dissociation along dorsal and ventral streams, underlies visual perception and action

**DOI:** 10.1101/609487

**Authors:** Dipanjan Ray, Nilambari Hajare, Dipanjan Roy, Arpan Banerjee

**Affiliations:** Cognitive Brain Dynamics Lab, National Brain Research Centre, Manesar, India 122052

**Keywords:** Dual Stream, Plasticity, Predictive Coding, Perception, Action

## Abstract

Visual dual stream theory posits that two distinct neural pathways of specific functional significance originate from primary visual areas and reach the inferior temporal (ventral) and posterior parietal areas (dorsal). However, there are several unresolved questions concerning the fundamental aspects of this theory. For example, is the functional dissociation between ventral and dorsal stream driven by features in input stimuli or is it driven by categorical differences between visuo-perceptual and visuo-motor functions? Is the dual stream rigid or flexible? What is the nature of the interactions between two streams? We addressed these questions using fMRI recordings on healthy human volunteers and employing stimuli and tasks that can tease out the divergence between visuo-perceptual and visuo-motor models of dual stream theory. fMRI scans were repeated after seven practice sessions that were conducted in a non-MRI environment to investigate the effects of neuroplasticity. Brain activation analysis supports an input-based functional dissociation and existence of context-dependent neuroplasticity in dual stream areas. Intriguingly, premotor cortex activation was observed in the position perception task and distributed deactivated regions were observed in all perception tasks thus, warranting a network level analysis. Dynamic causal modelling (DCM) analysis incorporating activated and deactivated brain areas during perception tasks indicates that the brain dynamics during visual perception and actions could be interpreted within the framework of predictive coding. Effectively, the network level findings point towards the existence of more intricate context-driven functional networks selective of “what” and “where” information rather than segregated streams of processing along ventral and dorsal brain regions.

## Introduction

The existence of two distinct streams of neural information processing-ventral and dorsal, projecting from the primary visual cortical areas to the inferior temporal and posterior parietal cortices, has been a powerful theory over almost 40 years (Mishkin and Ungerleider, 1982; Goodale and Milner, 1992). Similar duplex architectures in information processing associated with other brain functions e.g., auditory (Romanski et al., 1999), haptic (James and Kim, 2010), chemosensory (Frasnelli et al., 2012), attention (Vossel et al., 2014), speech (Hickok and Poeppel, 2007), and language (Saur et al., 2008) have been subsequently proposed. Despite such importance, several aspects of the visual dual stream theory are still poorly understood. Among them are specific roles of individual brain regions in the dual stream pathways, their interactions with each other for processing a putative perception/action task, and the functional reorganization of the engagement of dual stream regions with learning and familiarity. We address two prominent issues in the present article.

First, there exist diverging predictions regarding functional specialization of the two streams from the two most powerful variations of visual dual stream theory, the Mishkin-Ungerleider (MU) model (Mishkin and Ungerleider, 1982) and the Milner-Goodale (MG) model (Goodale and Milner, 1992. The MU model, also referred to as the visuo-perceptual model, suggests that the nature of input information determines the neural pathway for processing. Features that help in object identification (“what”) e. g., color, shape, texture etc. are processed in the occipito-temporal or ventral stream whereas perception of spatial (“where”) information e.g., position, velocity, depth, orientation are processed in the occipito-parietal or dorsal stream (Haxby et al., 1991; Horwitz et al., 1992; Haxby et al., 1994). In contrast, the MG model, alternatively referred to as the visuo-motor model, suggests that output or the task goal decides the selection of a processing pathway, ventral or dorsal. The ventral stream areas are needed for internal representation (“perception”) of both what and where information, whereas the dorsal stream areas are recruited for processing that same input information for guiding visuomotor “actions”. The MG model was supported by the observation that patient DF could insert a card in a slot, whose orientation was randomly set at different angles, relatively seamlessly, but was unable to correctly describe or otherwise report the orientation of the slot. While both the MG and MU models may predict same brain activation in certain situations as represented in Figure 1.a, e.g., ventral stream activation during “perception” of “what” information, nonetheless, the models diverge in activation prediction during situations such as “perception” of “where” information represented in Figure 1.b. Moreover, in some conditions e.g., during “action” task guided by “what” information represented in Figure 1.c, both the models anticipate activation in the same brain regions but the underlying pattern of interaction between different brain regions differs. The MU model was extensively supported by PET and fMRI studies and the MG model’s support comes predominantly from visual agnosia and optic ataxia studies (Schenk and Mcintosh, 2010). Thus, the first objective of the present work is to critically assess MU and MG models of visual dual stream in a single fMRI study.

**Figure 1.**
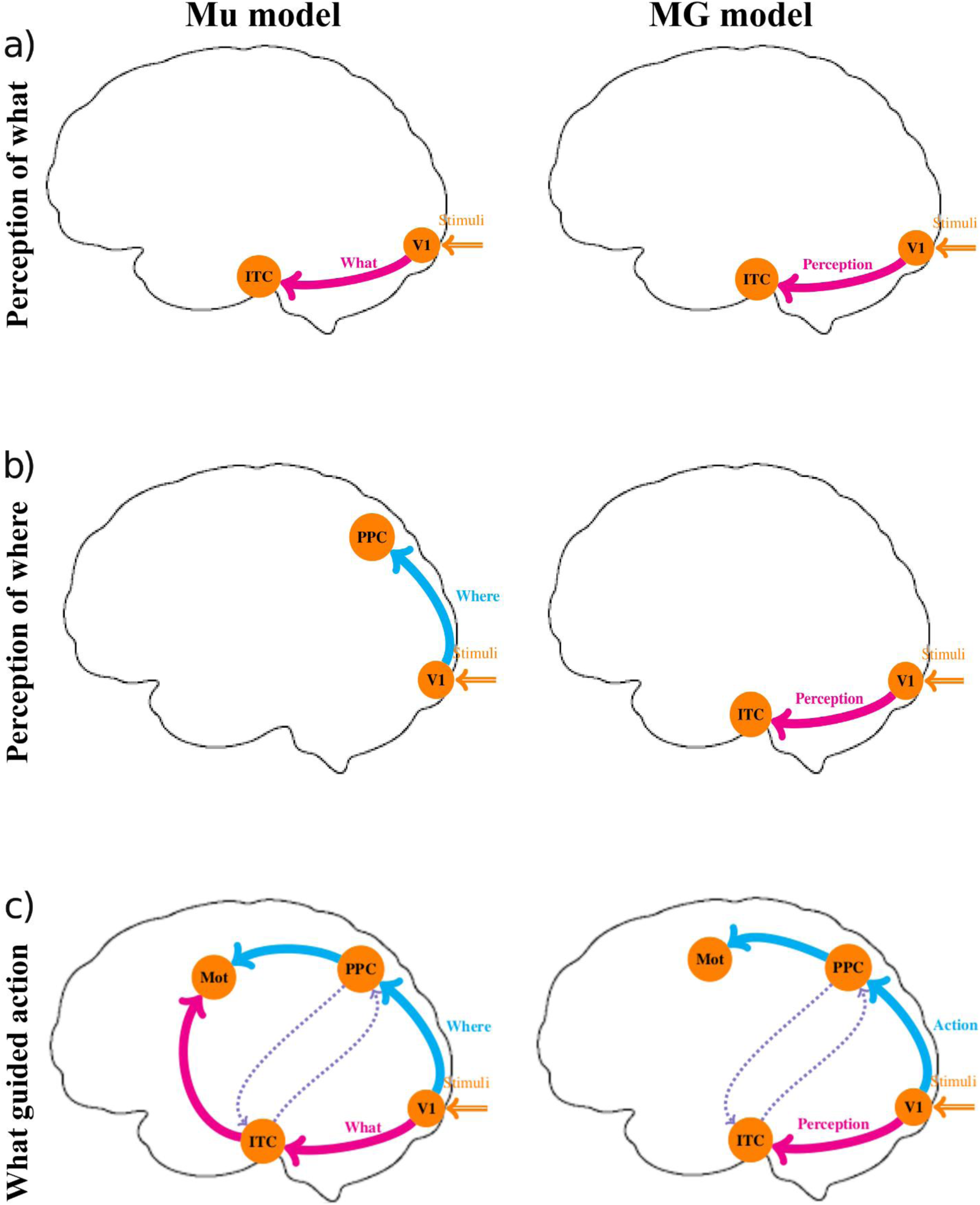
Predictions of brain activation by two models of dual stream theory in different tasks. (a) During perception of what information (e.g., color perception) both models predict activation of ventral stream regions but follow different rationale based on input and output level explanations, respectively. (b) During perception of where information (e.g., perception of position), however, the predictions of two models diverge. (c) For what information guided action tasks (e.g., reaching to a particular color target), though activation of primary visual areas, ventral and dorsal stream regions, and motor cortex is predicted by both the models, speculation about the flow of information between these regions is different. Particularly, according to the MG model, dorsal stream is independently capable of processing both what and where information for guiding action in the motor cortex. Thus, though there is the flow of information from the primary visual cortex to ventral stream regions, as there is the simultaneous internal representation of visual information while performing the action, and though the information from the ventral stream may feed into the dorsal stream, it is the dorsal stream that ultimately guides the action and the flow from ventral stream to motor cortex is redundant.

The second objective of the study is to probe into what extent the brain dynamics during visuo-perceptual and visuo-motor functions are subjected to reorganization by learning and familiarity. Longitudinal studies involving patients with visual form agnosia and optic ataxia resulting from ventral or dorsal stream damage have often yielded contradictory observations (Schenk, 2006; V.H. and KR., 2008; Schenk and Mcintosh, 2010; Schenk, 2012; Whitwell et al., 2015). For example, Schenk (2012) reported when haptic feedback was removed, DF was unable to insert the card in the target slot, that suggests a dissociation of action and perception is unlikely. Contrary to the Schenk (2012) study, Whitwell et al. (2015) reported that even with the removal of haptic feedback, DF was able to seamlessly insert the card in the target slot, essentially emphasizing the dominance of MG model. Since there was a period of 3 years that elapsed between these two studies, the effects of learning in the same patient DF cannot be ruled out while interpreting the contrary reports. Therefore, we hypothesized that parametric control of neuroplasticity can help in reconciling some of these apparently disparate observations. Moreover, if developmental changes are tracked, they can consequently be used to conceptualize a marker to differentiate between normal visuomotor functions and pathological scenarios. Hence, to explore the effects of neuroplasticity driven by behavioral skill development, we performed successive brain scans interspersed by a week of training in visual perception and action tasks.

## Materials and Methods

### Participants

Twenty-two right-handed healthy volunteers (14 females, 8 males) were included in the study who declared normal or corrected-to-normal vision with no history of neurological/ neuropsychiatric ailments. Two of the volunteer’s data were excluded due to excessive head movement inside the scanner. The mean age was 25.35 years (SD =2.796) in the final analysis. Handedness was tested according to the Edinburgh Inventory, to make sure participants were indeed right-handed. All participants gave written informed consent to the experimental procedure, the format of which was approved by the Institutional Human Ethics Committee of National Brain Research Centre (IHEC, NBRC) and in agreement with the Declaration at Helsinki.

### Experimental Design, Stimuli, and Tasks

An experimental paradigm was designed to reveal the brain activations along ventral and dorsal processing pathways in the context of attributes color, face, and position (Figure 2). Two kinds of tasks were conceptualized:

1. *Perception tasks (Color, face, position)*: Color perception was studied using four different-color filled circles that were presented one at a time randomly but successively, following which the participant was asked to report verbally the number of times the target color (red) was presented in a run.. Similarly, in face perception task, four different faces were randomly presented one at a time and the task was to indicate the number of times a particular target face was presented. In position perception trials, two black dots were presented in different positions with respect to the central cross in the screen, and the task was to calculate and report the number of times the two dots were equidistant from the central cross. Participants had to remember the number of times a target appeared for each category for the whole duration of the run. The number ranged from 0 to 4 (which varied randomly from task to task and across the sessions) to minimize the working memory load. In all three kinds of contexts, stimuli were presented at the center of the screen. In order to minimize eye movements and cued saccades during position perception tasks, the location of two black dots was restricted within the foveal vision (3 degrees of visual angle) of each participant. Visual angle extended by color stimuli and face stimuli were also 3 degrees. All stimuli were presented using Presentation software (Version 18.0, Neurobehavioral Systems, Inc., Berkeley, CA; www.neurobs.com).
2. *Visually-guided action tasks*: Participants were asked to move the cursor on the screen with the help of an fMRI compatible joystick (Current Designs, Inc.; Model HHSC-JOY-5; http://curdes.com) whose movement was calibrated to match the velocity and direction of the cursor movement to a target stimulus. Red circle, a specific target face, or the distant black dot from the center of the screen were the target stimuli among two dissimilar stimuli of the same category (color/face/position) presented simultaneously (Figure 2).

**Figure 2.**
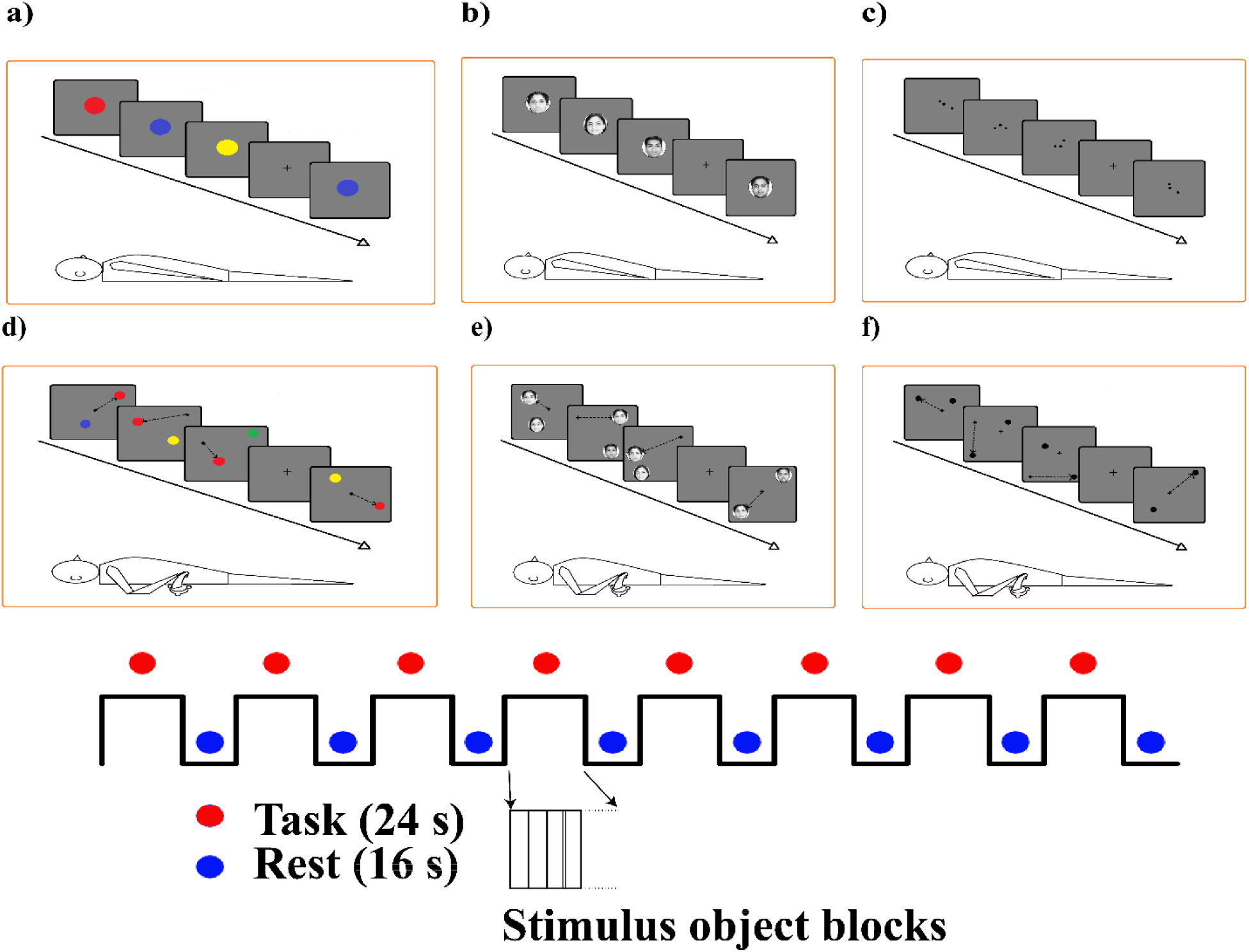
Experimental paradigm. In perception tasks, the participants were asked to calculate the number of times the target stimuli, (a)the red color filled circle, (b)the target face, (c) the equidistant black dots, were presented. In action tasks, they were instructed to move the cursor in the screen with the help of a joystick to the target stimuli, (a) the red dot, (b) the target face, (c) the distant black dot, among two simultaneously presented stimuli in each trial. Each trial was presented for 1 second in perception tasks and till the cursor reach the target (unless it is more than 4 seconds) in the action tasks. Stimuli were presented in active blocks of 24 seconds duration alternating with 16 seconds rest blocks. Each task run consists of such 16 alternating blocks. Presentation of the tasks for each participant and presentation of the stimuli within each block were randomized.

Visual stimuli for both perception and action tasks were presented with a grey background in eight “On” blocks (duration 24 seconds each) alternating with “Off” blocks of 16 seconds duration (Figure 2). During Off blocks, a central cross on a grey background was presented. 8 On and Off blocks (1 run) of each attribute were presented successively. In perception tasks, each stimulus was presented for one second with no inter-stimulus interval while in action tasks each stimulus persisted until the participants moved the cursor to the target location. Visuo-motor task was completed when the cursor reached the boundary of the desired target location, following which the current trial ended, and the next target appeared on the screen. However, if the participant had failed to move the cursor to the target within a window of 4s, the next set of visual objects would appear automatically. For perception tasks, the number of times a target attribute appeared was reported by participants verbally after the completion of 1 run. For action tasks, the number of times the stimulus appeared within each one block depended on the performance of the participants.

To assess the effects of learning, participants were trained in the aforementioned tasks for 7 consecutive days in a non-MRI environment following the first fMRI scan session. Each practice session comprised of the same six tasks identical to scanning sessions but the order of presentation of the individual stimulus within a task was randomized within each of the sessions. The order of six task blocks was also randomized. The number of practice sessions was decided based on a pilot study probing the improvement of response time with practice. From the eighth day of training the performance saturated in the pilot sessions.

### MRI Data Acquisition

Neuroimaging was performed with a 3T (Philips Achieva) magnetic resonance imaging (MRI) scanner at NBRC using a standard whole head coil (8-channels). To limit head movement-related artifacts, participants were verbally instructed to keep their heads as still as possible. Additionally, the participant’s head was cushioned by foam padding. Earplugs and customized noise-canceling headphones were used to attenuate scanner noise. The room lights were dimmed at near-identical levels for all participants.

Structural MRI: High-resolution T1-weighted structural MRI images with a repetition time (TR)= 8.4 ms, echo time (TE)=3.7ms, flip angle (FA) = 8 degrees, matrix = 252×230×170, field of view (FOV) = 250×230×170 mm were acquired from each participant for anatomical coregistration.

Functional MRI: T2* weighted functional whole-brain images were acquired with TR= 2000 ms, TE= 35 ms, FA = 90 degrees, matrix = 60×62×30, FOV = 230×230×179 mm during each task performance using a gradient echo-planar imaging (EPI) sequence.

### Behavioral Data Analysis

For perception tasks, the verbal response was sought from participants after each run to report the number of times the target stimulus was presented. For visually guided action tasks, the response time (RT) was defined by the time taken by the participant to move the cursor to the target object after two objects change position. Two-way ANOVA was employed to compare RTs across days and perception tasks as independent variables. A post-hoc Tukey-Kramer test was also used to compare RTs in all possible pairs of conditions.

### Pre-processing and brain activation mapping

The pre-processing and statistical analysis of fMRI data were executed with the SPM8 toolbox (Statistical Parametric Mapping, http://www.fil.ion.ucl.ac.uk/spm/). Initial 8 seconds of scanning sequence were discarded to allow the magnetization to stabilize to a steady state. Prior to statistical analysis, images were slice-time corrected, realigned with the mean image, motion corrected, coregistered with the corresponding T1-weighted images, normalized to a Montreal Neurological Institute (MNI, https://www.mcgill.ca/) reference template and resampled to 4×4×5 mm^3^. During motion correction, 2nd-degree B-Spline interpolation was used for estimation and 4th-degree B-Spline for reslicing. Coregistration used mutual information objective function while normalization used 4th-degree B-Spline interpolation. Temporal high pass filtering above 1/128 Hz was employed to remove low-frequency drifts caused by physiological and physical (scanner related) noises. Images were smoothed with a full-width at half-maximum (FWHM) Gaussian kernel 8×8×10 mm^3^.

The general linear model (GLM) based one-sample t-test was employed to identify brain activations and deactivations (Friston et al., 1994). The design matrix included regressors of interest for each task representing the event onsets and their time course as well as realignment parameters for head movement as regressors of no interest. The resulting statistical parametric maps of the t-statistics for the contrast “Task-Baseline” were thresholded at p < 0.01 (False Discovery Rate: FDR corrected) to get the activated voxels at each participant-level across the whole brain. Group analyses were performed using a random effects model. Deactivated voxels during tasks were identified by implementing a GLM with the contrast “Baseline-Task” and repeating the aforementioned steps. Anatomical localization of local maxima of activation/deactivation was assessed using the SPM Anatomy toolbox (v 2.2b, Eickhoff et al. 2005).

Subsequently, we were interested in tracking the number of activated/ deactivated voxels as well as the percentage signal change in dual stream areas between two scanning sessions interspersed with practice sessions. V1-V2 mask was created by combining BA17 and BA18 masks, ventral stream (VEN) mask by combining ventral extrastiate cortex, lateral occipital cortex, and fusiform gyrus and dorsal stream (DOR) mask by combining dorsal extrastiate cortex, V5/MT+, inferior parietal cortex, intraparietal sulcus, and superior parietal cortex. Probabilistic cytoarchitectonic maps from SPM Anatomy toolbox (Eickhoff et al., 2005) were used as masks for ROI computation. Comparison between the number of activated/deactivated voxels and percentage signal change in 2 scanning sessions was done using Wilcoxon signed rank test.

### Dynamic causal modelling

A deterministic bilinear variant of dynamic causal modelling (DCM) (Friston et al., 2003) was employed to probe the effective connectivity among the activated/deactivated regions. Alternative models were compared by Bayesian model selection, that rests on computing the model evidence, i.e., the probability of the data (BOLD signal) given a specific model. The posterior probability of coupling parameters were estimated by Bayesian Model Averaging (BMA), where we averaged over models, weighted by the posterior probability of each model. Effective network models were constructed for activation and deactivation separately in each hemisphere incorporating all ROIs. Different network schemas involving primary visual cortex (V1), ventral extrastriate areas (VES), fusiform gyrus (FG), dorsal extrastriate areas (DES), superior parietal lobule (SPL), pre-motor cortex (PMC), and motor cortex (Mot) as ROIs were chosen as nodes of “activation” (in perception and action tasks) and “deactivation” (in perception tasks) networks. There were no deactivated regions in action tasks (see “Mapping functional brain activity along dual stream: SPM results” in “Results” section).

Time series extraction: Time series for DCM analysis were extracted by taking the first principal component of the time series from all voxels included in a sphere of 6 mm diameter centered on the peak activated voxel in each participant. We also adjusted data for “Effects of interest” thus effectively mean-correcting the time series.

Model space construction: Plausible DCMs for activation networks in color and face perception tasks included bilateral intrinsic connectivity between primary visual cortex (V1) and extrastriatal ventral stream (VES), as well as between VES and fusiform gyrus (FG). No direct intrinsic connectivity between V1 and FG was considered. The recurrent or self-connections were also considered (Figure 3.a). Two kinds of model families were considered, in both of which visual inputs enter the system via primary visual cortex. However, in model 1, only the feed-forward connections, i.e., from V1 to VES and from VES to FG were modulated, whereas, in model 2, both feed-forward and feedback connections including from FG to VES and from VES to V1 were modulated.

**Figure 3.**
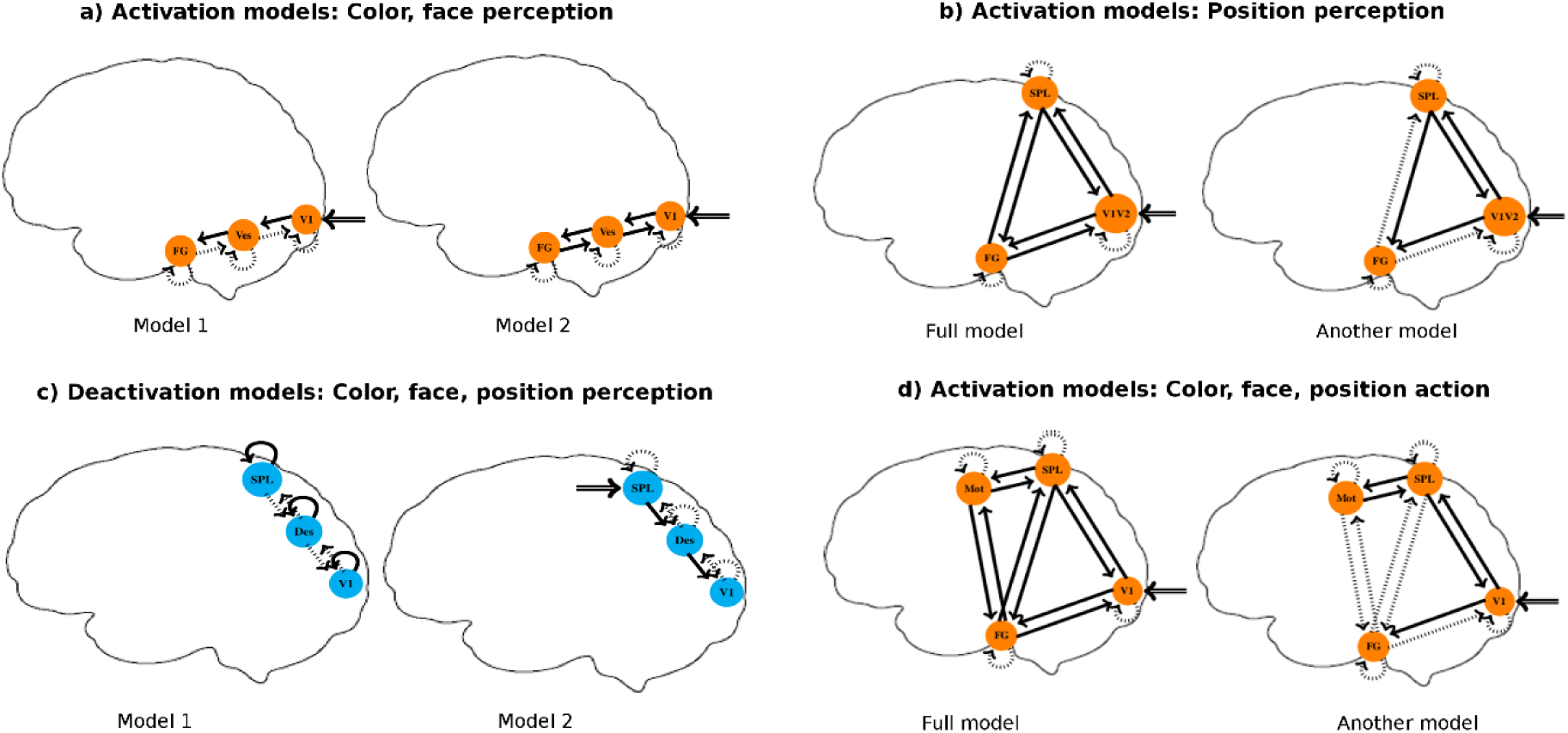
Model space. For (a) ventral stream activation in color and face perception, two competing models were compared. Primary visual cortex (V1), extrastriate ventral stream regions (Ves), and fusiform gyrus (FG) consisted the regions of interests (ROIs). For each model, intrinsic connections were assumed between ROIs in the immediate hierarchy, and self-loops. The visual inputs entered the network via the V1 node. In the first model, the modulation of only feedforward connections was considered while in the second model, in addition, the modulation of feedback connections was also considered. For (b) activation in position perception, we constructed and analyzed DCMs consisting of primary and secondary visual cortex (V1V2), fusiform gyrus (FG), and superior parietal lobule (SPL) ROIs. In all models, visual inputs entered the network via the V1V2 node. All the nodes were intrinsically connected among each other. The task-induced modulations of all non-self connections were considered. In total 27 DCMs were constructed based on the modulation of combinations of effective connections between three nodes. A “full” model and another arbitrarily chosen “alternative” model are shown in the figure. V1, extrastriate dorsal stream regions (DES), and superior parietal lobule (SPL) consisted the ROIs for (c) dorsal stream deactivation in color, face and position perception. Two competing models were compared. Like (a), for each model, intrinsic connections were assumed between ROIs in the immediate hierarchy, and self-loops. In the first model, modulation of self loops were considered and in the second model the modulation of feedback connections were considered. In model 2 visual inputs entered the network via the V1 node. For (d) activation in action tasks connections between V1, FG, SPL, and primary motor cortex (Mot) were considered. No direct intrinsic connection between V1 and Mot was considered, otherwise, all possible connections including self-loops were considered. All possible modulation of non-self connections gave rise to 80 competing models. For illustration, a full model with modulation of all non-self connections and another model with modulation of 5 connections were depicted.

Interestingly, significant voxel activations in primary visual cortex (V1) were not observed for position perception most likely owing to cognitive subtraction of proximate lower level mechanisms between task (central dots) and rest condition (central cross) both within 3 degrees of visual angle. However, significantly activated voxels could be found when we considered a combined ROI consisting of primary and secondary visual cortex (V1V2) that was created combining BA17 and BA18 regions from Talairach Daemon database (http://www.talairach.org/daemon.html). Hence, we constructed and analyzed DCMs consisting of V1V2, FG, and superior parietal lobule (SPL) ROIs for position perception task (Figure 3.b). FG and SPL ROIs were same as those used in action tasks. In all models, visual inputs entered the network via the V1V2 node. All the nodes were intrinsically connected among each other. The task-induced modulations of all non-self connections were considered. In total 27 DCMs were constructed based on the modulation of combinations of effective connections between three nodes. A “full” model and another arbitrarily chosen “alternative” model are shown in Figure 3.b.

DCMs for deactivation networks (during perception tasks) had bidirectional intrinsic connectivity among nodes in the immediate hierarchy (V1-DES and DES-SPL) and self-connections simultaneously (Figure 3.c). Out of the two models tested, model 1 had only the self-connections modulated whereas in model 2, input entered the system via SPL and all other top-down connections (SPL→ DES, DES→VIS) were modulated by the tasks.

Only activation networks were relevant for action tasks and models consisting of four ROIs - V1, FG (ventral stream area), SPL (dorsal stream area) and motor cortex (Figure 3.d) were considered. In all models, visual inputs entered the network via primary visual cortex. All the nodes were intrinsically connected among each other except primary visual cortex and motor cortex between which there was no direct intrinsic connection. The modulation of all “non-self” connections between nodes was considered. A “full” model in which all “non-self” connections were modulated is represented in Figure 3.d. Other models were constructed based on the modulation of combinations of effective connections between four nodes. One such model with modulation of 5 connections is also shown in the same figure. In total, 80 models were evaluated for model evidence computation.

## Results

### Behavioral performance and effects of practice

All participants were 100% accurate in counting the number of target stimuli that were presented in each block during perception tasks, during both scanning sessions and for the 7 practice sessions. Response times (RT) were computed trial-by-trial in visually guided action tasks (Figure 4). Two-way ANOVA on RT with task category (color, face, or position action) and training days as variable showed significant main effect of both practice days (F(8,486) = 32, p < 0.0001) and task condition (F(2,486) = 46.68, p < 0.0001) on RT. The interaction between these variables was not significant ( F(16,486) = 0.83, p = 0.5004). In general, color action showed the fastest and position action the slowest response time. Compared to the last practice session, RT deteriorated in 2^nd^ fMRI scan. Post-hoc analysis with the Bonferroni multiple-comparison test revealed that RT in 2^nd^ fMRI scan session is significantly faster than RT in 1^st^ fMRI scan session (p=0.0029).

**Figure 4.**
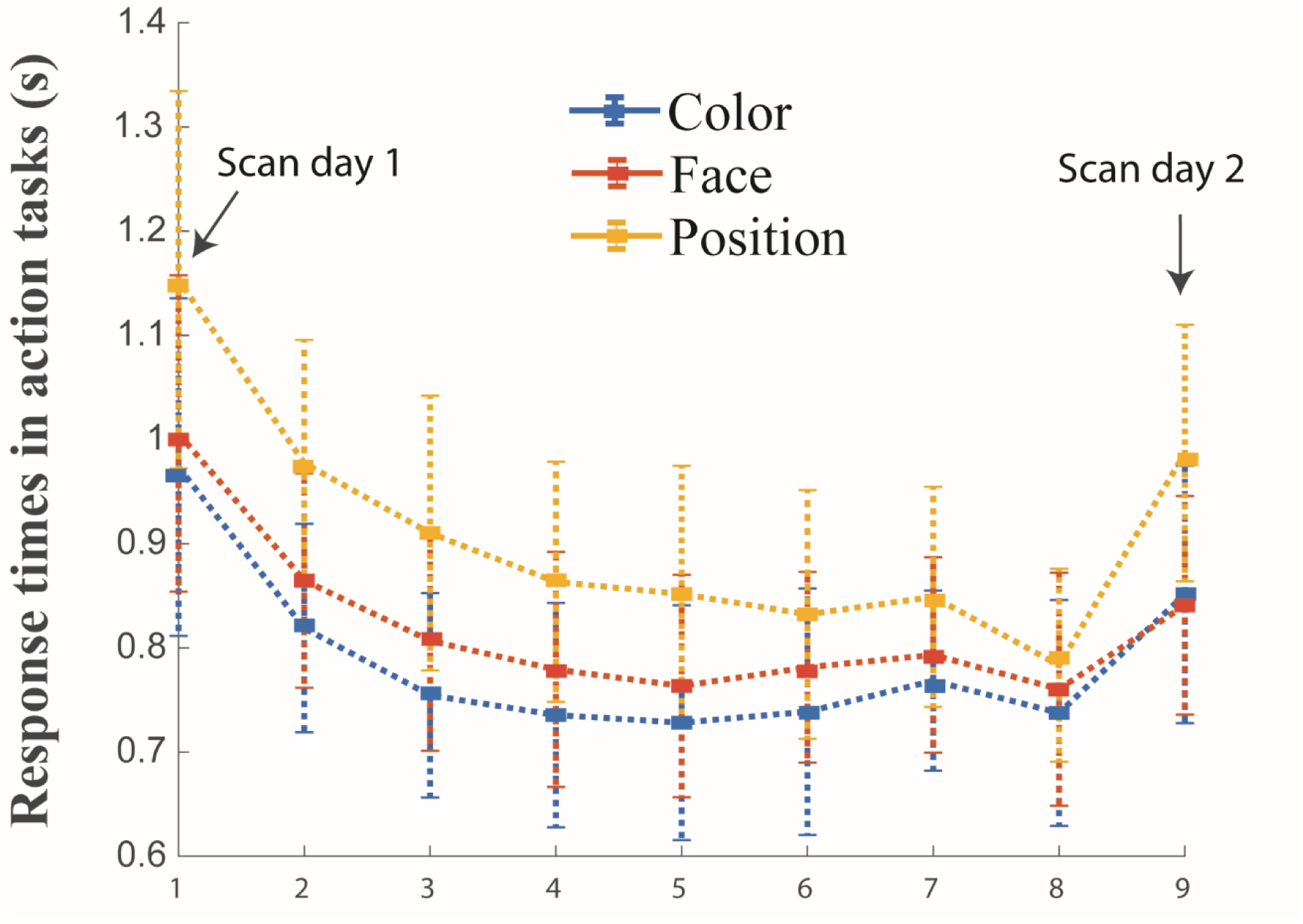
Behavioral result. Group level Mean and SD of response times (RTs) in color, face, and position action tasks across different sessions.

### Mapping functional brain activity along dual stream: SPM results

#### Activation and deactivation of dorsal and ventral visual areas in perception tasks

Significant activations were observed along primary visual areas (V1 and V2) and along the ventral stream (e.g., V3v, V4v, lateral occipital complex (LOC), and fusiform gyrus (FG)) during color and face perception tasks for both scanning sessions, separated by 7 practice days (Figure 5, Table 1).

**Figure 5.**
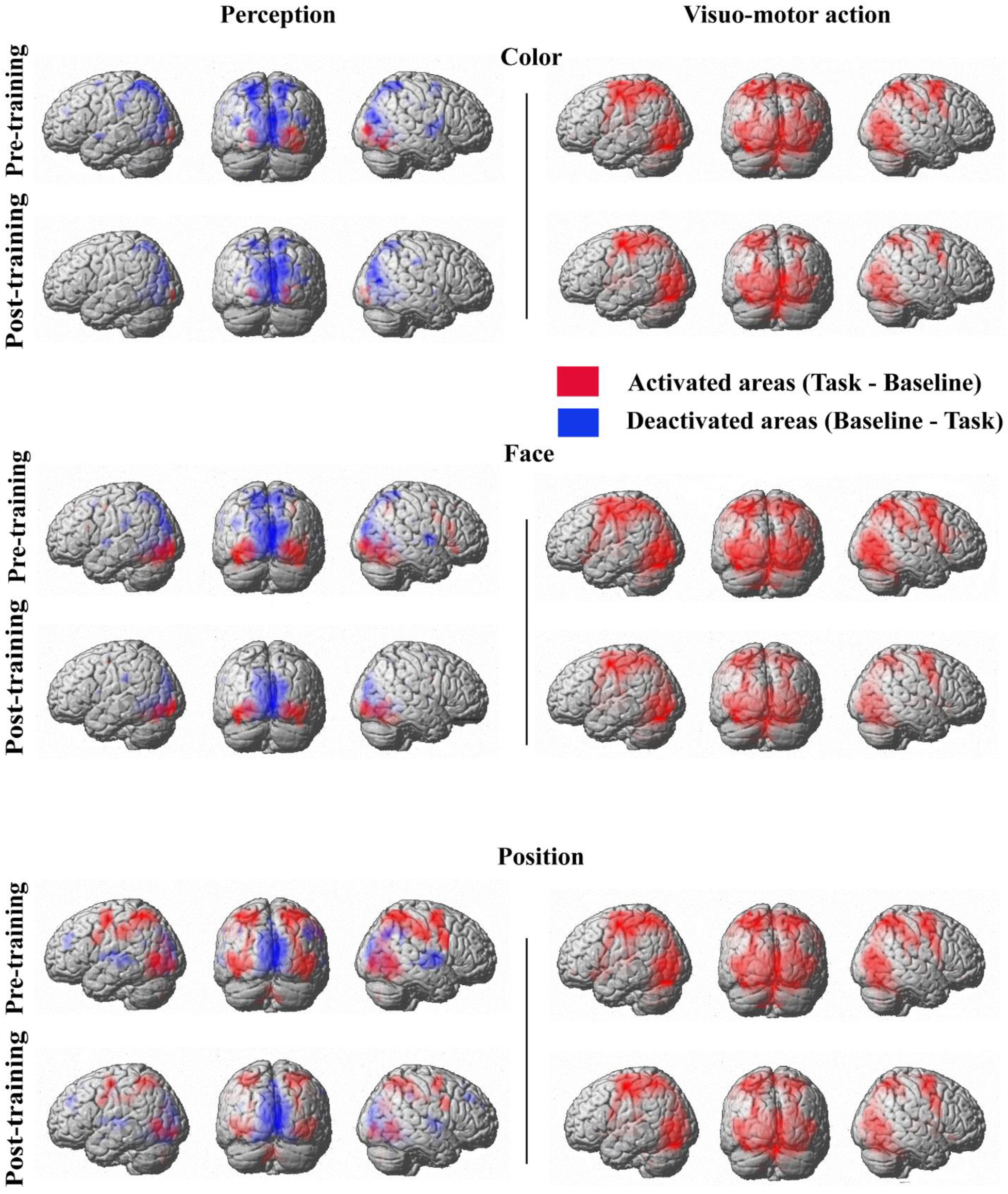
Group level brain activation and deactivation in perception and action tasks (pre- and post-training). Red and blue voxels represent activation and deactivation respectively in task blocks compared to rest blocks. For details of brain regions see Table 1.

**Table 1.**
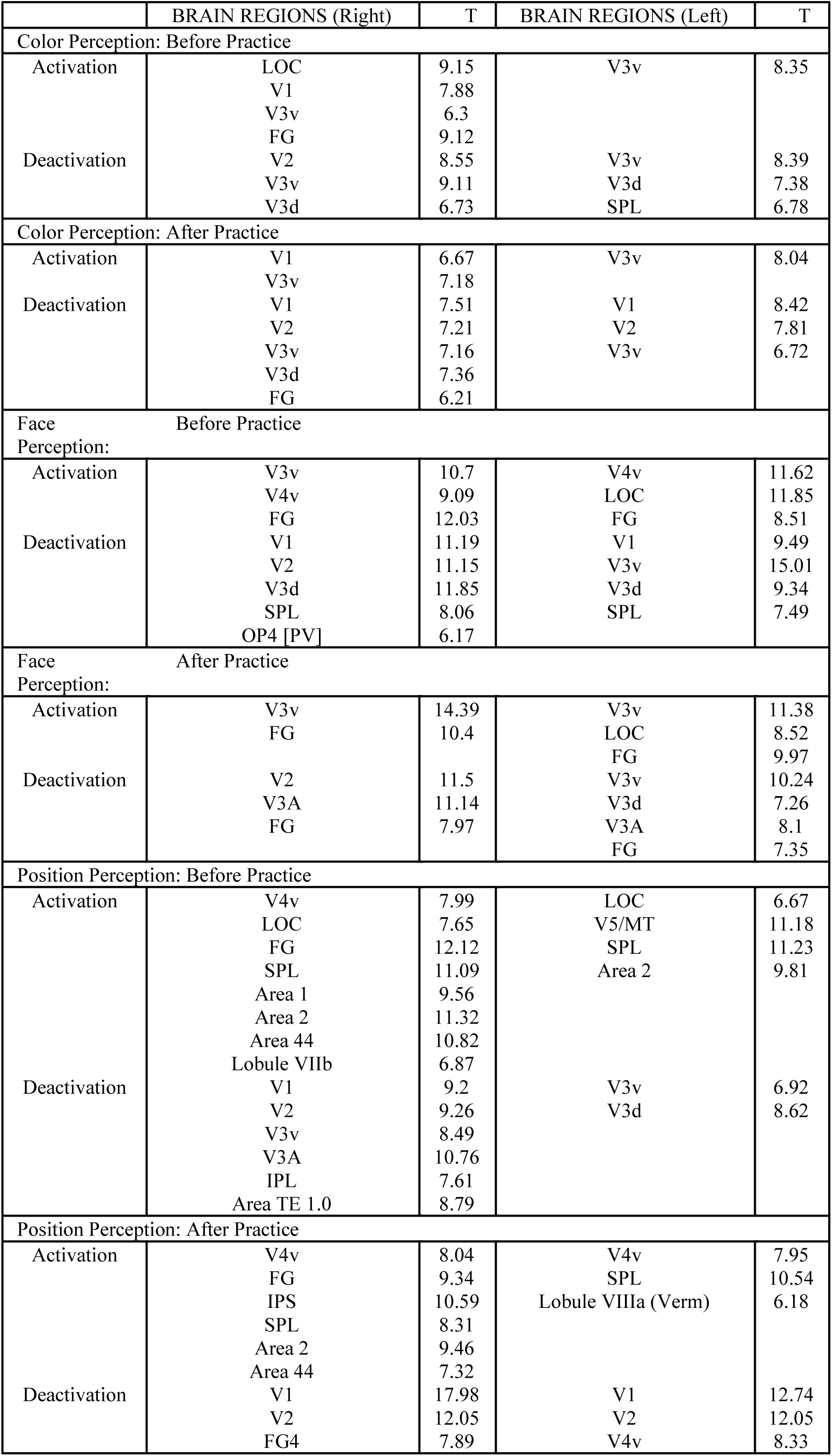
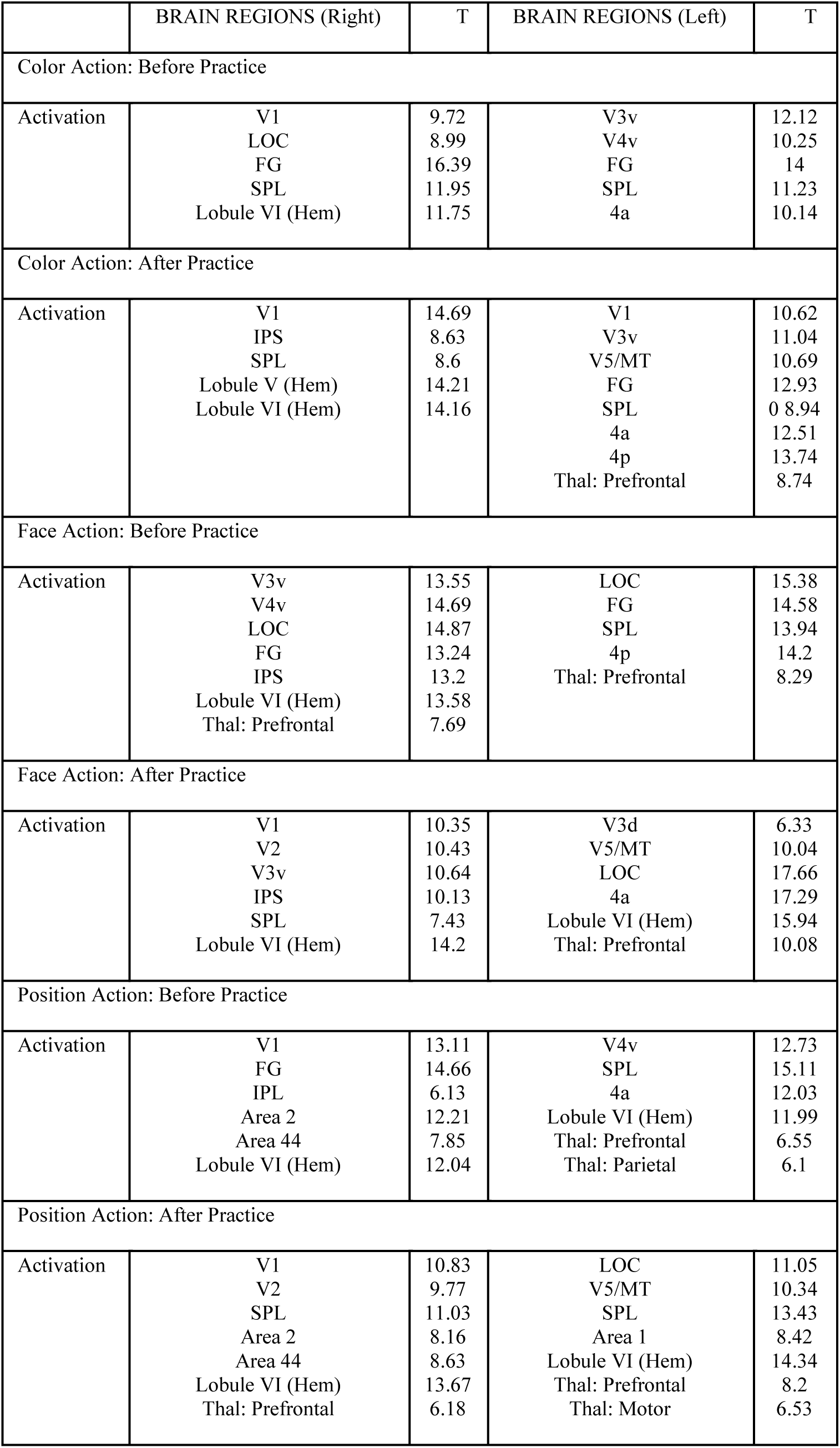
Brain regions showing local maxima of BOLD activity for Group Level Analysis and their corresponding T values. Brain regions were defined according to probabilistic cytoarchitectonic mapping used in SPM anatomy toolbox. v-ventral extrastriate cortex, d-dorsal extrastriate cortex, LOC-lateral occipital cortex, FG-fusiform gyrus, IPL-inferior parietal lobule, IPS-intraparietal sulcus, SPL-superior parietal lobule, OP-parietal operculum, TE 1.0-primary auditory cortex, Thal-thalamus, Lobule-cerebellar lobule.

In position perception task (Figure 5, Table 1), bilateral ventral (e.g., V4v, LOC, FG), and dorsal (e.g., V5/MT, SPL) stream regions were activated for both scan 1 and 2. Bilateral premotor cortex (PMC) also showed activation in both the scans. Interestingly, primary and secondary visual cortices did not exhibit activation in either scan at the false discovery rate (FDR)-corrected group level analysis.

Subsequently, the outcome of Wilcoxon signed ranked tests performed to evaluate the number of activated/ deactivated voxels change between scan 1 & scan 2 were reported in Figure 6.a-f. Similarly, results from Wilcoxon signed ranked tests performed on the percentage signal change comparisons between scan 1 and scan 2 were presented in Figure 6. g-l. A general trend of decrease in the extent of activated voxels in the ventral and dorsal stream areas in all perception tasks emerged from comparisons between scan 1 and 2. However, the percentage signal change between scanning sessions rarely changed.

**Figure 6.**
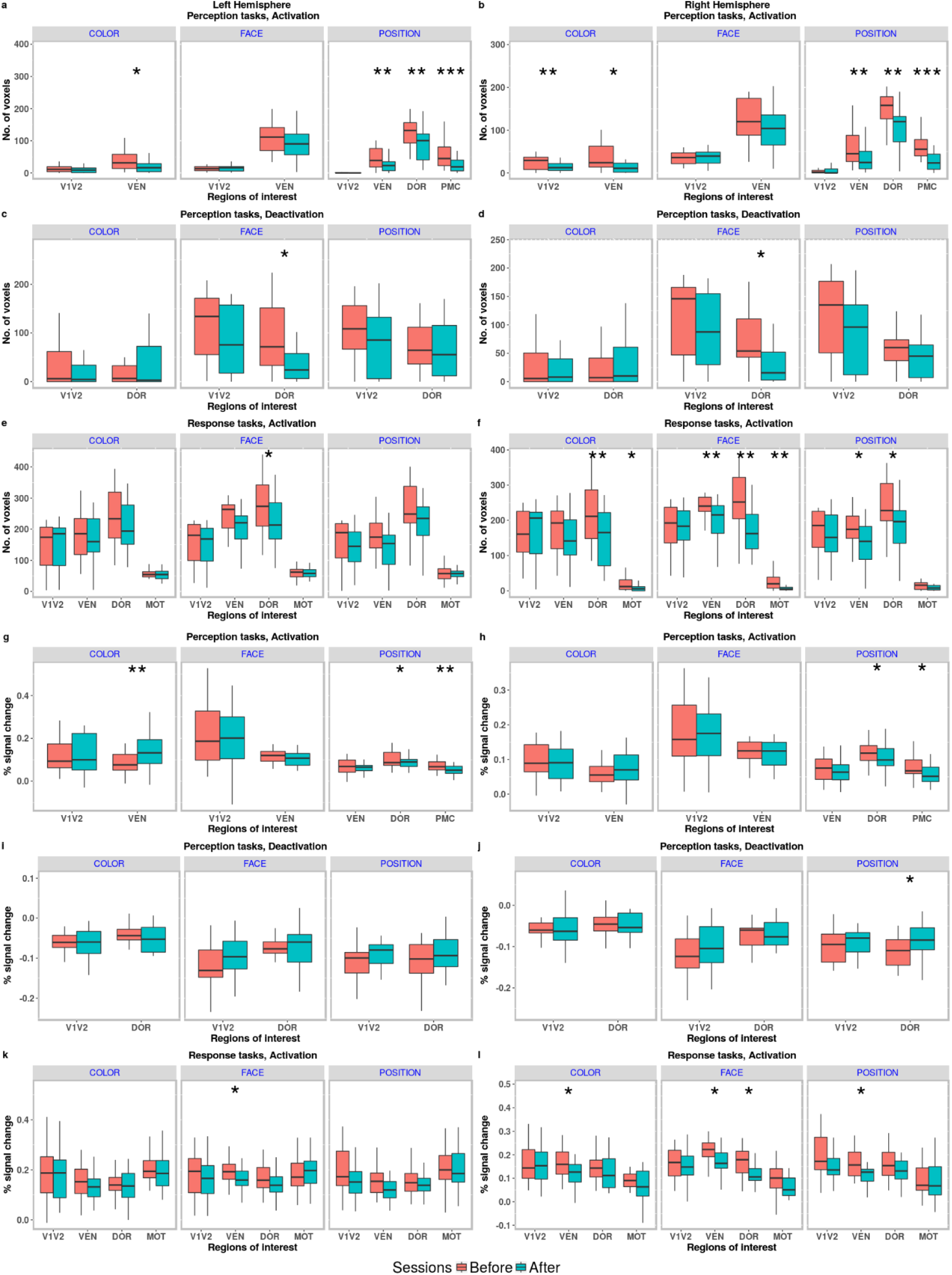
(a-f) No of voxels activated/deactivated during Perception and Response tasks (Before and after practice sessions). (g-l) Percentage signal change during Perception and Response tasks (Before and after practice sessions). V1V2-primary and secondary visual cortex, VEN-ventral stream, DOR-dorsal stream, PMC-premotor cortex, MOT-primary motor cortex. Stars above the boxplots indicate a statistically significant difference in no. of activated voxels/percentage signal change before and after practice. (Wilcoxon signed rank test).

Intriguingly, all perception tasks showed distinct areas of deactivation at the group level (Figure 5, Table 1). The deactivated areas predominantly involved bilateral primary and secondary visual cortices and dorsal stream regions such as extrastriate dorsal stream and superior parietal lobule (SPL). Certain ventral stream regions, particularly extrastriate ventral stream region, also showed deactivation during certain perception tasks. Compared to activated areas in perception (and response) tasks, the deactivated areas were located more medially. In contrast to activation, there was no statistically significant change in the extent of deactivation between scan 1 and scan 2, except dorsal stream deactivation in face perception (Figure 6. a-d).

#### Activation of dorsal and ventral stream areas in action tasks

In all action tasks (Figure 5, Table 1) primary and secondary visual cortices, ventral and dorsal stream areas and motor cortex underwent bilateral activation in both scan 1 and 2. There was a decrease in the extent of activations, however, the statistically significant decrease during scan 2 predominantly occurred in the right hemisphere (Figure 6. e-f). Analogous to perception tasks, percentage signal change did not show significant change with practice in scan 2 (Figure 6.k-l) compared to scan 1. Unlike perception tasks, there were no significant deactivations in any of the action tasks, in both scan 1 and 2.

### Brain network analysis

After identifying the activation and deactivation of several brain regions in perception and action tasks, we tried to underpin the effective connectivity among these regions across tasks, and their alteration with practice. To address these systematically, we employ dynamic causal modelling (DCM) to evaluate effective brain networks underlying perception and action tasks according to the scenarios proposed in Figure 3.

### Perception tasks

#### Effective connectivity among activated regions

Model 1 in color and face perception tasks captures only feed-forward sensory driven processing circuit for networks involving activated brain regions (Figure 3). On the other hand, model 2 addressed a network schema that involves both feed-forward and feed-back processing. For activation networks during color and face perception, model 2 schemas were more likely candidates that facilitate the underlying information processing. Subsequently, on parameter estimation (see Figure 7.a), all the feed-forward connections among activated regions were found to be positive whereas feedback connections were negative. Scan 1 and scan 2 had a similar strength of effective connections.

**Figure 7.**
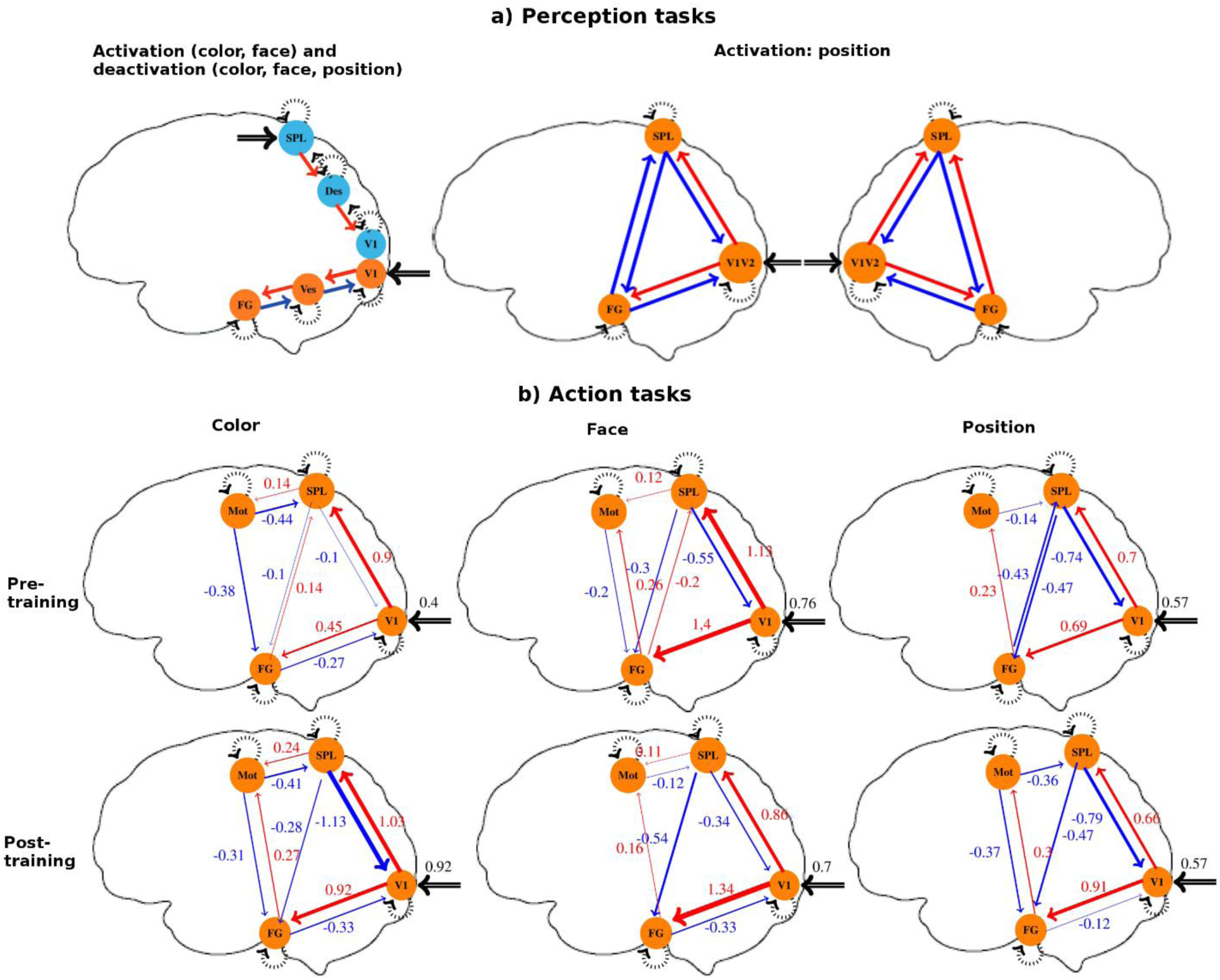
Effective connectivity using dynamic causal modelling (DCM) during perception and action tasks. (a) Perception tasks: The winning model configuration for ventral stream activation and dorsal stream deactivation during color and face perception tasks, and activation in primary and secondary visual cortex (V1V2), ventral (FG), and dorsal (SPL) stream regions during position perception tasks (positive modulation: red, negative modulation: blue). Pre-and post-training analyses show similar values of estimated coupling parameters for each connection for color and face perception. (b) Action tasks: The winning model configuration for pre- and post-training sessions. The color of the arrow represents the nature (positive modulation: red, negative modulation: blue), and the thickness of the arrow represents the mean value of the estimated coupling parameter for modulation of effective connectivity. V1-primary visual cortex, VES-extrastriate ventral stream regions, FG-fusiform gyrus, DES-extrastriate dorsal stream regions, SPL-superior parietal lobule, Mot-primary motor cortex.

In position perception tasks all possible combinations of connection schemas among V1V2, SPL and FG were considered (Figure 3.b). The mean coupling parameters indicated that feedforward connections between V1V2 and nodes of ventral (FG) and dorsal (SPL) streams are excitatory whereas feedback connections are inhibitory (Figure 7.a, right panel). There was a trend of V1V2 to SPL coupling being slightly higher than V1V2 to FG coupling (Table 2), the differences however were not significant. There were also strong inhibitory influences from SPL to FG bilaterally across both the sessions which did not show any significant change with practice. FG to SPL influences, on the other hand, were inhibitory in left hemisphere but excitatory in right hemisphere in both sessions.

**Table 2.**
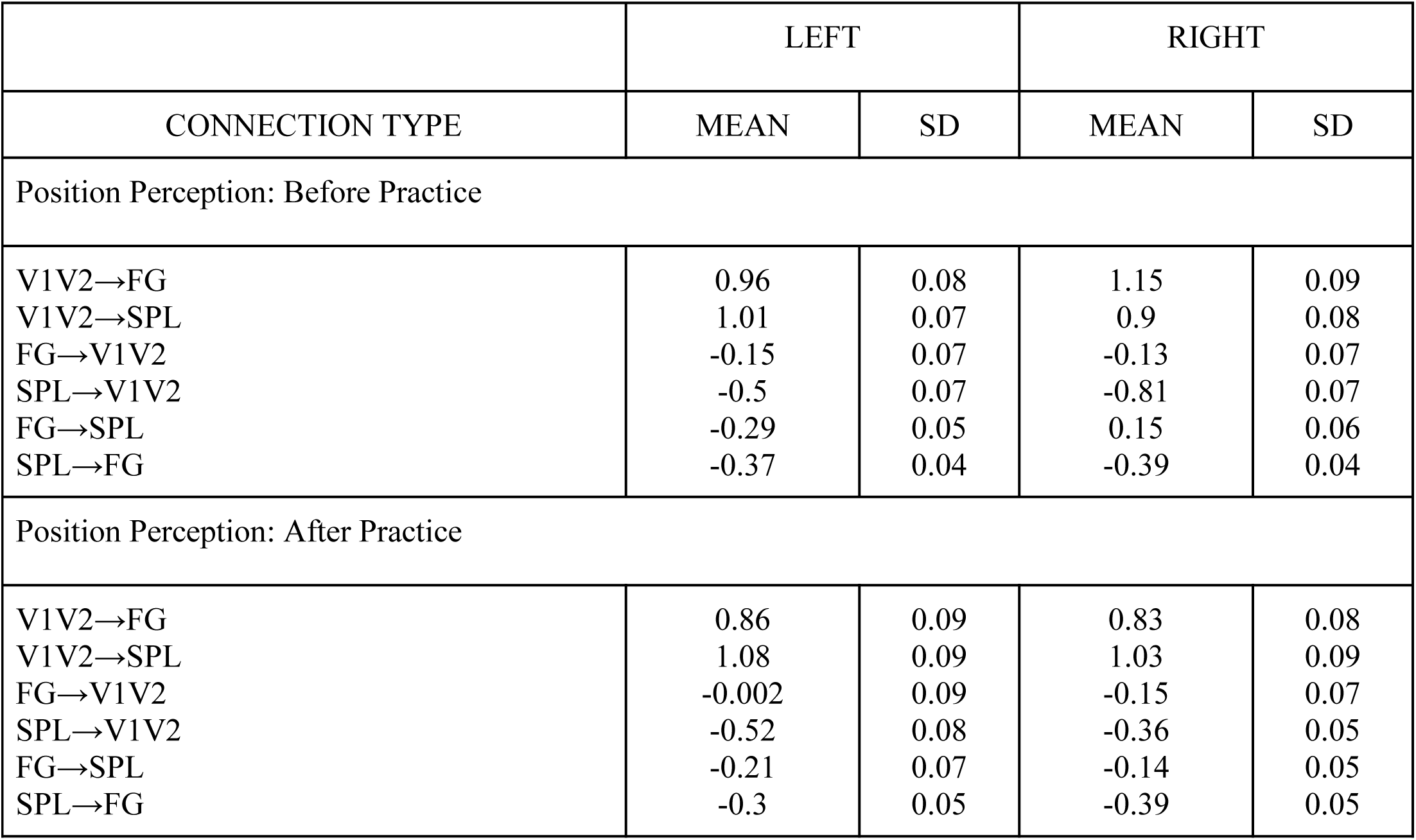
DCM parameter estimates of modulatory connections during position perception task. V1V2 - primary and secondary visual cortex, FG - fusiform gyrus, SPL - superior parietal lobule.

#### Effective connectivity among deactivated brain regions

Connection schemas with only self-connections within nodes (model 1) and with feedback-connections between nodes with input at SPL (model 2) were tested (Figure 3.c). Model 2 schemas were favored and the input to SPL was found to be inhibitory whereas the coupling parameters for feedback connections between deactivated regions were estimated to be positively modulated during each perception tasks for both scanning session 1 and 2.

#### Effective connectivity among brain regions during action tasks

DCM analysis of action tasks required comparison among 80 different models (Figure 3.d). On estimating the coupling parameters, we found that primary visual cortex positively influenced ventral and dorsal regions as predicted by dual stream theory in all action tasks (see Figure 7. b). Both the ventral region (FG) and dorsal region (SPL), in turn, positively influenced the motor cortex to perform the visually guided action tasks cued with face and color stimuli. In position action tasks, the motor cortex was driven by FG but not SPL. The feedback connections (FG→V1, SPL →V1, MOT→FG, MOT→SPL), when present, were all inhibitory. There was also strong inhibitory influence from dorsal stream regions to ventral stream regions while performing movements and this inhibitory influence either remained same (for position action task) or was enhanced (for color and face action) with practice as reflected in the estimated coupling parameters of DCM.

## Discussion

Our study aimed to investigate the visual dual stream theories proposed by Mishkin-Ungerleider (MU) (Mishkin and Ungerleider, 1982; Mishkin et al., 1983) and Milner-Goodale (MG) (Goodale and Milner, 1992; Milner et al., 2012), on a task ideally designed to validate their respective predictive power. Accordingly, we conceptualized two kinds of tasks - one that involved the perception of visual objects (perception tasks), e.g., color, face or position stimuli in absence of any motor goal and the other that required performance of goal-directed movements (action tasks) with a joy-stick following color, face or position cues. MU model would predict only dorsal stream activations for position stimuli but ventral stream activations for color and face stimuli in perception tasks. On the other hand, MG model would predict the involvement of only ventral areas in all perception tasks. Intriguingly, we saw both dorsal and ventral stream activations in position perception tasks, an observation that diverged from predictions from either of the models. However, identification of an object is implicit in localizing its position, thus the observations were not completely inconsistent with the input-based MU model. Secondly, we observed patterned deactivation in dorsal stream brain regions for color/face and position stimuli. Thirdly, the activation and deactivation in perception and action tasks showed changes in the pattern depending on the context of the tasks (e.g., the hand used in performing the action). Fourthly, using dynamic causal modelling (DCM) (Friston et al., 2003), we could demonstrate how predictive coding may be a more plausible network-centric model for understanding the role of top-down modulations of lower- and higher-order visual areas during perception-action tasks. DCM also revealed how “cross-stream” inhibitory influences are exerted by dorsal stream regions onto ventral stream areas during action and position perception tasks, which upon training, either remained same or got consolidated to a unidirectional dorsal to ventral influence. Recently, increasing evidence has shown that the ventral and dorsal streams are not strictly independent, but do interact with each other directly (for a review see van Polanen and Davare (2015)). However, this is the first study, to the best of our knowledge, to point out that the nature of dorsal to ventral influence may be inhibitory and demonstrate the evolution of such interactions with training. Based on all these observations, we propose a revision of stream-based models to a more nuanced network-level understanding of visual information processing that emphasizes functional integration as a binding principle of visual processing in perception/action contexts.

BOLD deactivation is relatively a rarely discussed topic and often looked upon with suspicion by the neuroimaging community. More often than not it is explained by the so-called “blood stealing” effect - redirection of blood flow to the activated region and away from adjacent inactive regions, and routinely ignored (Wade, 2002; Hayes and Huxtable, 2012). Nonetheless, the deactivation found during the perception tasks in the present study was consistent across tasks and practice sessions, was much more extensive compared to the activation (at least in color and face perception) and included too many distal regions than the activated areas to share a common pool of blood supply. Thus, neuronal suppression is a more physiologically grounded explanation for the deactivations we observed in this study in contrast to blood stealing (Frankenstein et al., 2003).

The decrease in the number of activated voxels in perception and action tasks with practice reflected the habituation effect, a form of neuroplasticity marked by the progressive decrease of the responses to repeated sensory stimulation (Glaser and Whittow, 1953). In action tasks, the lateralization of the shrinkage of activated regions denoted that the habituation in the dual stream is dependent on the context e.g., right-handedness of the participants in the present study. Preservation of overall activation pattern, the constancy of percentage signal change in the face of shrinkage and lowering of RT, supported the idea that habituation effectuates more efficient processing of information which consumes a lesser amount of energy reflected by a decrease in the spatial boundaries of activation patterns (Kok et al., 2012).

The predictive coding framework, an emerging theory of brain function, suggests that the brain is continually attempting to predict the external causes of sensory information at all levels of the cortical processing hierarchy (Mumford, 1992; Rao and Ballard, 1999; Friston and Kiebel, 2009). According to the most recent variation (Friston and Kiebel, 2009) of this view, feedback connections from a higher- to a lower-order sensory cortical area carry predictions of lower-level neural activities and inhibit/explain away the predicted signal in the lower level. The residual error, if any, is carried by the feed-forward connections, that are excitatory in nature, and that update the prediction at the higher level. This process continues until prediction matches the incoming stimulus. This view represents a more computationally efficient alternative to the traditional model of sensory processing where each feature of the sensory object is processed and integrated in a predominantly bottom-up schema. The present results indicated feed-forward connections among activated regions in perception tasks to be contributing towards excitatory “influences” while feedback connections contributing to inhibitory “influences” (see Figure 7.a), thus complying with the variation of predictive coding theory proposed by Friston and Kiebel (2009).

The current results also reflected neural suppression in the dorsal stream in perception tasks to be mediated by top-down inhibitory influence. A possible explanation of deactivation in the dorsal stream is repetition/expectation suppression (RS or ES) (Meyer and Olson, 2011; Grill-Spector et al., 2006), as in all perception tasks stimuli were presented centrally in the same location. As the stimuli location was fully predictable, there was no feedforward prediction error. On the other hand, as the subject concentrated to perceive the stimuli, the top-down inhibitory influence of prediction increased during active blocks, thus resulting in an overall deactivation compared to rest blocks. Thus, explanation of RS / ES based on predictive coding was supported by the top-down inhibitory influence revealed by DCM analysis. This explanation contradicts a more traditional explanation centered on local mechanisms such as fatigue (Grill-Spector et al., 2006) that can be represented by self-inhibiting loops to a neuronal population so that the inhibition is proportional to the neuronal activity (DCM 2 in our analysis).

The DCM analysis revealed a consistent inhibitory influence of SPL to FG during action and position perception tasks. There are already a few papers emphasizing the interaction between ventral and dorsal stream regions during task performance (Himmelbach and Karnath, 2005; van Polanen and Davare, 2015). However, to our knowledge, this was the first work to point out the nature of dorsal to ventral influence to be inhibitory.

The most important takeaway from our study is the urgent requirement of conceptualization of visual brain as a conglomeration of context-driven networks perpetually involved in integrative information processing (see also Schenk and Mcintosh, 2010) as opposed to two functionally dichotomous independent streams of processing. Most of the earlier works on dual stream emphasize on functional dissociation along distinct neural pathways with a scope of cross-talks while our work places functional integration mechanisms, not limited to cross-stream interactions but also between feedforward and feedback controls, at the core of a visual processing theory. Furthermore, our study emphasizes the role of inhibitory influences in shaping up visual information processing. Interestingly, the strengthening of the inhibitory influence over practice corresponds to the improvement of the response time in action tasks. To ascertain the exact role of this inhibitory influence, and the reason behind its strengthening would be exciting questions for future research. Electrophysiological study (including micro-electrode recordings from primate) could provide insight into the neurophysiological basis of the inhibitory influence by exploring the temporality of ventral and dorsal stream activity. Transcranial magnetic stimulation (TMS) study could be explored as an alternative approach in human participants. Specific brain regions in the ventral or dorsal stream could be stimulated while performing visuomotor tasks and their effect on the behavior (response time, accuracy) could be studied in the near future.

In conclusion, our study establishes that both dual-stream models of information processing cannot sufficiently capture the patterns of functional brain activations and connectivity, although the importance of input signal characteristics remain paramount. The connectivity results specifically showed that visual information is processed by a highly interconnected cortical network in which lateral and feedback connections among lower and higher order visual areas play very important roles. The particular pattern that both dorsal areas like SPL and ventral areas like FG inhibits primary visual areas which in turn have positive (excitatory) influences on the dorsal and ventral areas but never vice-versa is a strong support for the mechanisms of predictive coding to be operational in visual processing. This calls for a shift from the present stream-based models to a more nuanced network-level understanding of visual processing that can address the aspects of information integration vis-à-vis predictive coding hypothesis.

## Author contributions

DR1 (Ray) and AB conceived the study; DR1, NH, and AB collected the data; DR1, NH, DR2 (Roy) and AB analyzed the data; DR1, DR2, and AB wrote the manuscript.

## Data availability statement

The data that support the findings of this study are available from the corresponding author upon request for academic purposes. However, the data remains the property of National Brain Research Centre an autonomous institute of Department of Biotechnology, Government of India.

## Acknowledgments

We thank Dr. Moumita Das for her help in data analysis, Dr. Soibam Shyamchand Singh for his helpful comments on improving the readability of the manuscript, NBRC Core funds, and infrastructural support. DR (Ray) was supported by a Cognitive Science Research Initiative Fellowship (SR/CSRI/PDF-13/2014) from Department of Science and Technology, DR (Roy) was supported by the Ramalingaswami fellowship (BT/RLF/Re-entry/07/2014) and DST-CSRI extramural grant (SR/CSRI/21/2016) and AB was supported by Ramalingaswami fellowship BT/RLF/Re-entry/31/2011) and Innovative Young Biotechnologist Award (IYBA), (BT/07/IYBA/2013).

## Notes

**Conflict of interest statement:** The authors declare no competing financial interests.

## References

Eickhoff, S. B., Stephan, K. E., Mohlberg, H., Grefkes, C., Fink, G. R., Amunts, K., and Zilles, K. (2005). A new SPM toolbox for combining probabilistic cytoarchitectonic maps and functional imaging data. NeuroImage, 25(4):1325–1335.

Frankenstein, U., Wennerberg, A., Richter, W., Bernstein, C., Morden, D., Remy, F., and Mcintyre, M. (2003). Activation and deactivation in blood oxygenation level dependent functional magnetic resonance imaging. Concepts in Magnetic Resonance, 16A(1):63–70.

Frasnelli, J., Lundstrom, J. N., Schopf, V., Negoias, S., Hummel, T., and Lepore, F. (2012). Dual processing streams in chemosensory perception. Frontiers in human neuroscience, 6:288.

Friston, K. and Kiebel, S. (2009). Predictive coding under the free-energy principle. Philosophical Transactions of the Royal Society B: Biological Sciences, 364(1521):1211–1221.

Friston, K. J., Harrison, L., and Penny, W. (2003). Dynamic causal modelling. NeuroImage, 19(4):1273–302.

Friston, K. J., Holmes, A. P., Worsley, K. J., Poline, J.-P., Frith, C. D., and Frackowiak, R. S. J. (1994). Statistical parametric maps in functional imaging: A general linear approach. Human Brain Mapping, 2(4):189–210.

Glaser, E. M. and Whittow, G. C. (1953). Evidence for a non-specific mechanism of habituation. The Journal of physiology, 122(Suppl):43–4P.

Goodale, M. a. and Milner, A. (1992). Separate visual pathways for perception and action. Trends in Neurosciences, 15(1):20–25.

Grill-Spector, K., Henson, R., and Martin, A. (2006). Repetition and the brain: neural models of stimulus-specific effects. Trends in Cognitive Sciences, 10(1):14–23.

Haxby, J. V., Grady, C. L., Horwitz, B., Ungerleider, L. G., Mishkin, M., Carson, R. E., Her-scovitch, P., Schapiro, M. B., and Rapoport, S. I. (1991). Dissociation of object and spatial visual processing pathways in human extrastriate cortex. Proceedings of the National Academy of Sciences of the United States of America, 88:1621–1625.

Haxby, J. V., Horwitz, B., Ungerleider, L. G., Maisog, J. M., Pietrini, P., and Grady, C. L. (1994). The functional organization of human extrastriate cortex: a PET-rCBF study of selective attention to faces and locations. The Journal of Neuroscience, 14:6336–6353.

Hayes, D. J. and Huxtable, A. G. (2012). Interpreting deactivations in neuroimaging. Frontiers in psychology, 3:27.

Hickok, G. and Poeppel, D. (2007). The cortical organization of speech processing. Nature reviews. Neuroscience, 8:393–402.

Himmelbach, M. and Karnath, H.-O. (2005). Dorsal and Ventral Stream Interaction: Contributions from Optic Ataxia. Journal of Cognitive Neuroscience, 17(4):632–640.

Horwitz, B., Grady, C. L., Haxby, J. V., Schapiro, M. B., Rapoport, S. I., Ungerleider, L. G., and Mishkin, M. (1992). Functional associations among human posterior extrastriate brain regions during object and spatial vision. Journal of cognitive neuroscience, 4:311–322.

James, T. W. and Kim, S. (2010). Dorsal and Ventral Cortical Pathways for Visuo-haptic Shape Integration Revealed Using fMRI. In Multisensory Object Perception in the Primate Brain, pages 231–250. Springer New York, New York, NY.

Kok, P., Jehee, J., and deLange, F. (2012). Less Is More: Expectation Sharpens Representations in the Primary Visual Cortex. Neuron, 75(2):265–270.

Meyer, T. and Olson, C. R. (2011). Statistical learning of visual transitions in monkey inferotemporal cortex. Proceedings of the National Academy of Sciences, 108(48):19401–19406.

Milner, A. D., Ganel, T., and Goodale, M. A. (2012). Does grasping in patient d.f. depend on vision? Trends in Cognitive Sciences, 16(5):256–257.

Mishkin, M. and Ungerleider, L. G. (1982). Contribution of striate inputs to the visuospatial functions of parieto-preoccipital cortex in monkeys. Behavioural Brain Research, 6(1):57–77.

Mishkin, M., Ungerleider, L. G., and Macko, K. A. (1983). Object vision and spatial vision: two cortical pathways. Trends in Neurosciences, 6:414–417.

Mumford, D. (1992). On the computational architecture of the neocortex. II. The role of cortico-cortical loops. Biological cybernetics, 66(3):241–51.

Rao, R. P. N. and Ballard, D. H. (1999). Predictive coding in the visual cortex: a functional interpretation of some extra-classical receptive-field effects. Nature Neuroscience, 2(1):79–87.

Romanski, L. M., Tian, B., Fritz, J., Mishkin, M., Goldman-Rakic, P. S., and Rauschecker, J. P. (1999). Dual streams of auditory afferents target multiple domains in the primate prefrontal cortex. Nature neuroscience, 2(12):1131–6.

Saur, D., Kreher, B. W., Schnell, S., Kummerer, D., Kellmeyer, P., Vry, M.-S., Umarova, R., Musso, M., Glauche, V., Abel, S., Huber, W., Rijntjes, M., Hennig, J., and Weiller, C. (2008).Ventral and dorsal pathways for language. Proceedings of the National Academy of Sciences, 105(46):18035–18040.

Schenk, T. (2006). An allocentric rather than perceptual deficit in patient D.F. Nature neuroscience, 9(11):1369–70.

Schenk, T. (2012). No dissociation between perception and action in patient DF when haptic feedback is withdrawn. The Journal of neuroscience, 32(6):2013–7.

Schenk, T. and Mcintosh, R. D. (2010). Do we have independent visual streams for perception and action? Cogn neurosci., 44(0):1–44.

van Polanen, V. and Davare, M. (2015). Interactions between dorsal and ventral streams for controlling skilled grasp. Neuropsychologia, 79(Pt B):186–91.

V.H., F. and KR., G. (2008). Grasping visual illusions: consistent data and no dissociation. Cogn Neuropsychol., 25(7-8):920–50.

Vossel, S., Geng, J. J., and Fink, G. R. (2014). Dorsal and ventral attention systems: distinct neural circuits but collaborative roles. The Neuroscientist, 20(2):150–9.

Wade, A. R. (2002). The Negative BOLD Signal Unmasked. Neuron, 36(6):993–995.

Whitwell, R. L., Milner, A. D., Cavina-Pratesi, C., Barat, M., and Goodale, M. A. (2015). Patient df’s visual brain in action: Visual feedforward control in visual form agnosia. Vision Research, 110(Pt B):265–276.

